# DISSECT: DISentangle SharablE ConTent for Multimodal Integration and Crosswise-mapping

**DOI:** 10.1101/2020.09.04.283234

**Authors:** Geoffrey Schau, Erik Burlingame, Young Hwan Chang

## Abstract

Deep learning systems have emerged as powerful mechanisms for learning domain translation models. However, in many cases, complete information in one domain is assumed to be necessary for sufficient cross-domain prediction. In this work, we motivate a formal justification for domain-specific information separation in a simple linear case and illustrate that a self-supervised approach enables domain translation between data domains while filtering out domain-specific data features. We introduce a novel approach to identify domain-specific information from sets of unpaired measurements in complementary data domains by considering a deep learning cross-domain autoencoder architecture designed to learn shared latent representations of data while enabling domain translation. We introduce an orthogonal gate block designed to enforce orthogonality of input feature sets by explicitly removing non-sharable information specific to each domain and illustrate separability of domain-specific information on a toy dataset.

## I. Introduction

The task of domain translation broadly relates to identifying a pair or set of transfer functions that model a data point in one domain as an equivalent data point in some other domain. For many problems, data may be collected from two complementary modalities for the same instance or event, enabling familiar statistical approaches for inferring correlative or causal relationships between measurements made in two or more domains. However, in instances for which paired measurements are not available, mapping a representation of a measurement into another domain becomes a significantly more complex challenge.

These types of situations arise often in the field of biomedical research in which cellular assays inherently destroy or permanently modifies the sample under study such that no two assays can be applied to the same sample. For example, the recent emergence of single-cell genomic sequencing technologies enable molecular characterization of individual cells, but destroy those cells in the process, therefore their application to the same set of cells is generally mutually exclusive. Similarly, single-cell multiplexed imaging technologies enable deep spatial profiling of tissue, yet the permanent fixture of tissue limits the applicability of sequencing technologies following or preceding imaging. In these cases, unpaired single-cell resolution measurements are available, though the absence of paired measurements limits the degree to which biological insights may be translated between technological approaches.

In this work, we consider the case of unpaired domain translation. Presently, literature describing learning systems for domain translation are largely applied to image-to-image translation settings. CycleGAN, built upon the generative adversarial framework [1], introduced the notion of cycle consistency to enable image-to-image domain translation in the absence of paired data [2], [3]. Although highly effective in certain settings, the authors recognize limitations of a purely GAN-based system, which experience common issues such as mode collapse [4]. Unlike GANs, variational autoencoders (VAEs) [5] learn optimal latent representations of input data by learning to generate reconstructions of an input data source rather than an arbitrary image capable of fooling a learned discriminator. Split-brain autoencoders [6] have recently been proposed as an alternative approach for learning cross-channel information of individual data samples, though the architecture has yet to be extended to the challenge of unpaired data samples from complementary domains. Shared latent representations between domains has been proposed using a pair of VAE models in both UNIT and MUNIT learning architectures [7], [8], though neither approach seeks to disentangle shared factors of variation.

In this paper, we consider the domain translation task between independent data domains that each describe the same underlying unit of analysis but from distinct, yet not strictly orthogonal, projections while estimating the degree to which complementary domains inform each other. First, we consider a simple linear case and illustrate the benefit of identifying domain-specific information. Second, motivated by theoretical derivation from a linear case, we propose a self-supervised deep learning-based approach that leverages unpaired data across two domains to learn crosswise mapping while disentangling mutual information content from distinct profiling modalities at single-cell resolution on a toy dataset.

## II. Problem Formulation

We consider a multimodal dataset where *X*_1_ and *X*_2_ are measurements from two different domains. For given *X*_1_ ∈ ℝ^*m*^ and *X*_2_ ∈ ℝ^*n*^, we are interested in identifying a crosswise mapping Φ(·), Ψ(·) as shown in Figure 1.

**Fig. 1.**
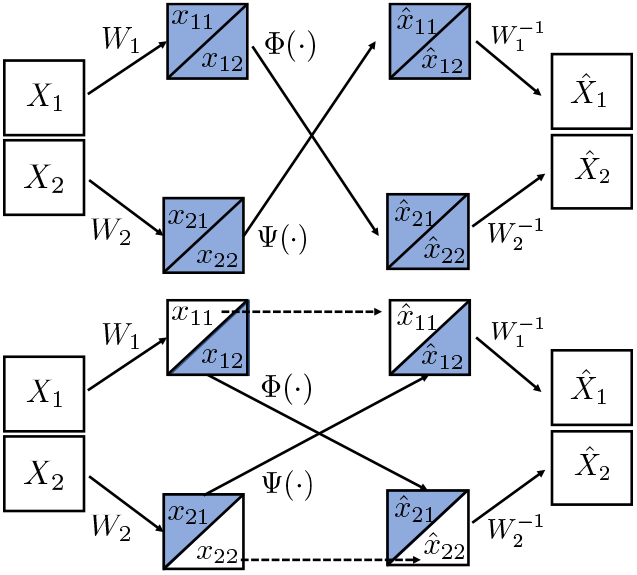
A conceptual diagram of crosswise mapping: (top) crosswise mapping without dissecting domain-specific features (similar to split-brain autoencoder) (bottom) Φ(·), Ψ(·) are any mappings (including autoencoder) where solid line represents shared network and dot line represent non-shared network.

**Fig. 2.**
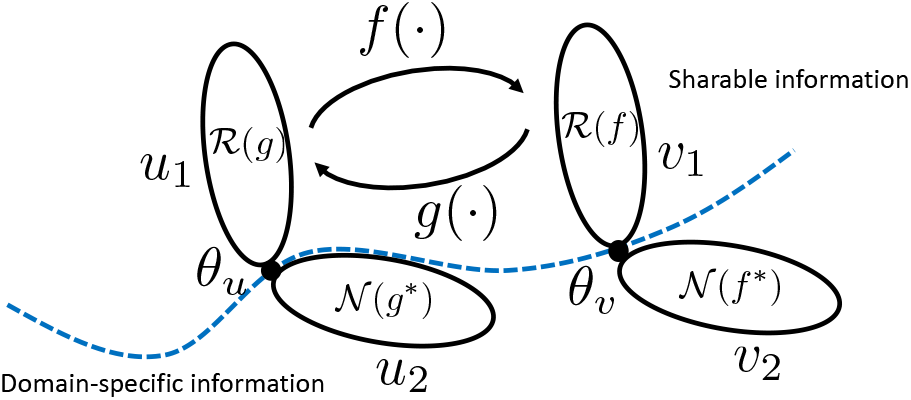
A continuous linear mapping case where crosswise mappings *f* : *u*_1_ → *υ*_1_ and *g* : *υ*_1_ → *u*_1_ are adjoint to each other.

### A. Disentangling domain-specific and sharable features to identify a crosswise mapping

In each domain, there exist domain-specific (i.e., *x_ii_*, *x_jj_*), and sharable contents/features (i.e., *x_ij_*, *x_ji_*) and we assume that they are disjoint, i.e., *x_ii_* ⊥ *x_ij_* or *x_ji_*. To split domain-specific and sharable content, we introduce a simple transformation *W* and assume that it is an orthonormal matrix (i.e., *W*^⊺^*W* = *I* and inner product (·, ·), (*w_i_*, *w_j_*) = 0 for *i* ≠ *j* and (*w_i_*, *w_j_*) = 1 for *i* = *j*) where *W*_1_ ∈ ℝ^*m*×*m*^ and *W*_2_ ∈ ℝ^*n*×*n*^ are orthogonal matrices:

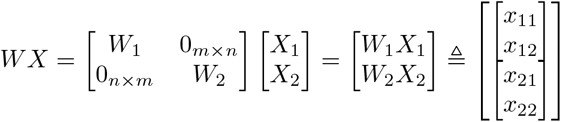

One could consider that *W* as a data sorting transform where each row of *W* consists of zeros and 1, i.e., standard basis, where corresponding nonzero component is chosen for either domain-specific or shared data set. Also, we consider the inverse mapping in a similar way:

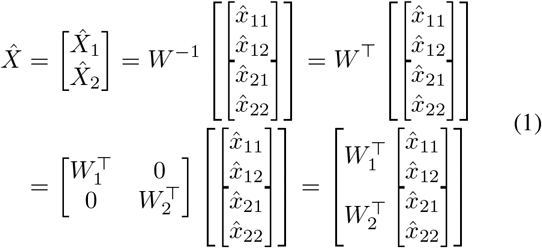

where 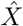 represents the reconstructed data from input data *X* with a crosswise mapping (Φ(·), Ψ(·)) as shown in Figure 1.

#### Lemma 1

*W* transformation does not affect the two-norm, i.e., 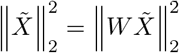

*Proof:* by definition, i.e., satisfies orthogonal invariance.

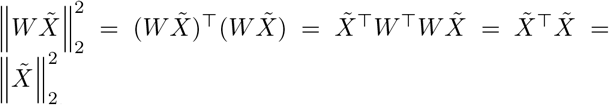

To learn crosswise mapping across two domains, first we need to identify sharable content and split them into two disjoint models or networks as shown in Figure 1 (bottom). To identify crosswise mapping (solid arrow), it is trained to perform prediction on one subset of data from another subset, similar to split-brain autoencoder [6]. For non-shared model (dotted arrow), we basically bypass them (mapping is an identify matrix or a simple autoencoder, i.e., input is the same as output). Note that if we do not consider disentanglement of sharable and domain-specific information (Figure 1 (top)), it is difficult to learn crosswise-mapping due to the non-sharable information across two domains.

#### Lemma 2

An orthonormal transformation *W* reduces reconstruction error 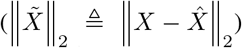 by bypassing disjoint information to non-shared network.

*Proof:* With *W* transformation, assume that *φ* and *ψ* are the optimal crosswise mappings to minimize the reconstruction error:

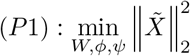

Then, with two disjoint networks, the reconstruction error is as follow:

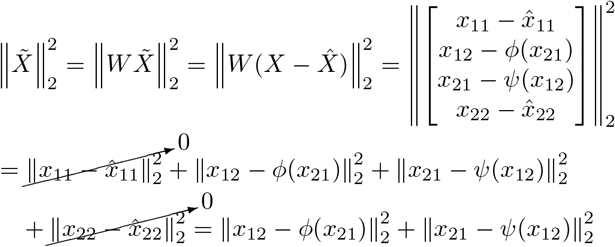

Without splitting into two disjoint networks (i.e., no bypass term), assume that Φ = (*ϕ*_1_, *ϕ*_2_) and Ψ = (*ψ*_1_, *ψ*_2_) are the optimal mappings to minimize the reconstruction error:

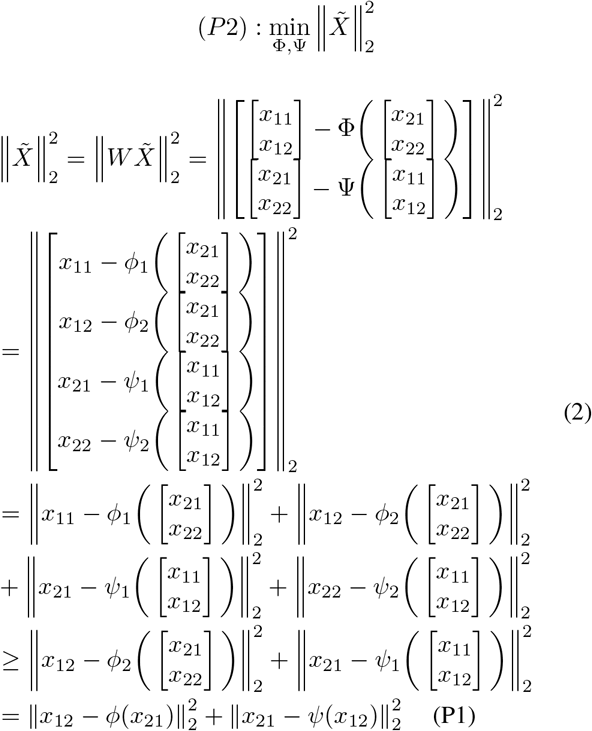

In the last equality sign, since we assume that *x*_22_ and *x*_11_ are disjoint from *x*_12_ and *x*_21_ (i.e., domain specific content) by definition, we can ignore these components.

Lemma 2 shows that (P1) provides the lower reconstruction error compared to (P2). Note that in Equation (2) (before the inequality sign), the first and last terms can cause huge reconstruction error since *x*_11_ and *x*_22_ are domain-specific features by definition and *x_ii_* and *x_ij_* (or *x_ji_*) are disjoint by the assumption, and thus this can be infinite value (i.e., no crosswisemapping from domain *i* to domain *j*). On the other hand, *W* allows us to dissect domain-specific features (*x*_11_, *x*_22_) and identify the shared (*x*_12_, *x*_21_) information across multimodal datasets by using two disjoint networks, rather than attempting to force highly divergent datasets into a completely shared latent space mapping (P2).

One may argue that (P1) may bypass the shared information to reduce the reconstruction error, for example, making *x*_12_ and *x*_21_ are empty set since we do not penalize any reconstruction error in non-shared part. To address this question, we consider the following case. First, we denote 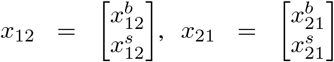 where superscript {·}^*b*^ and {·}^*s*^ represent the shared feature which go through “bypass” and “shared” networks respectively. Second, we assume that there exists 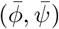 such that 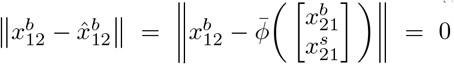 and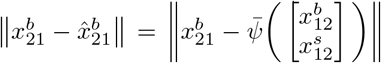, i.e., there exist a perfect crosswise mapping.

If we bypass this shared information, the reconstruction error can be written as follows: 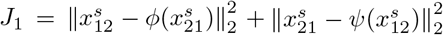. If we do not bypass this shared information, the reconstruction error can be written as follows:

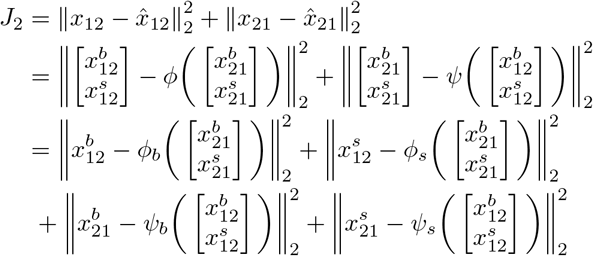

where we denote *ϕ* = (*ϕ_b_*, *ϕ_s_*) and *ψ* = (*ψ_b_*, *ψ_s_*).

#### Lemma 3

If there exists a crosswise mapping 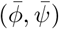 such that it guarantees zero or negligible reconstruction error (i.e., exact crosswise mapping), then bypassing those shared contents will be suboptimal, i.e., a larger reconstruction error, *J*_1_ ≥ *J*_2_.

*Proof:*

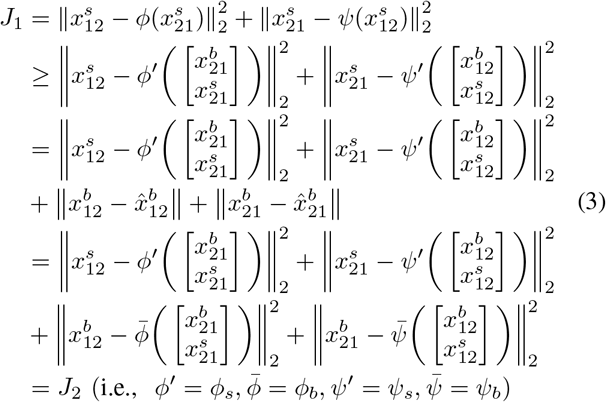

In the first inequality, we consider a mapping *ϕ′, ψ′* with keeping bypass term 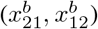 and without loss of generality, this mapping provides a lower error bound. Therefore, bypassing the shared information will result in the suboptimal solution where the shared information is necessary for reducing the reconstruction error in shared network. By Lemma 3, the optimization problem (P1) will identify *W* which only allows to bypass non-shared content information.

### B. Cross-domain cycle consistency to deal with unpaired dataset

The work presented herein leverages unpaired data across two domains to learn crosswise mapping while disentangling mutual information content from distinct profiling modalities. To address the challenge of unpaired data samples from complementary domains, we consider a cycle-consistency concept as free supervisory signal for learning a crosswise mapping across two domains. If we identify the exact crosswise mapping for the identified shared information, both Φ(·) and Ψ(·) satisfy Φ(Ψ(·)) = *I_m_*(·) and Ψ(Φ(·)) = *I_n_*(·) where Φ : ℝ^*n*^ → ℝ^*m*^ and Ψ : ℝ^*m*^ → ℝ^*n*^. Here we consider a simple linear case in order to build a theoretical framework to address questions of why we need disentanglement and cross-domain consistency for building deep learning models.

### Example: a linear mapping case

Consider *u* = *u*_1_ ⊕ *u*_2_ ∈ ℝ^*m*^ and *υ* = *υ_1_* ⊕ *υ*_2_ ∈ ℝ^*n*^ where subscript {·}_1_ represents sharable content and subscript {·}_2_ represent domain specific information, and ϕ represents the direct sum. Define continuous linear maps *f* : *u*_1_ → *υ*_1_ and *g* : *υ*_1_ → *u*_1_ and consider cyclic consistency loss:

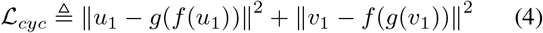

#### Proposition 1

*u*_1_ = *g*(*υ*_1_) and *υ*_1_ = *f*(*u*_1_) is the optimal solution for minimizing cyclic consistency loss (4).

*Proof:*

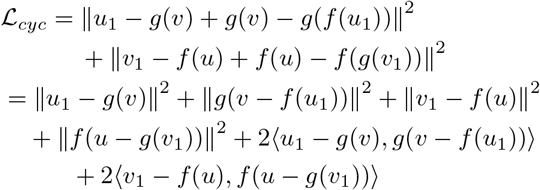

Plug in *u* = *u*_1_ + *u*_2_ = *g*(*υ*_1_) + *u*_2_ and *υ* = *υ*_1_ + *υ*_2_ = *f* (*u*_1_) + *υ*_2_, then

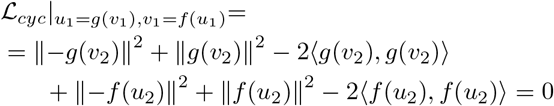

#### Remark 1.

Since 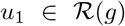 and *u*_2_ is the orthogonal complement in ℝ^*m*^, 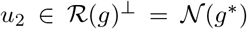 by definition where 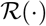 represents the range space, 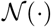 represents the null space, and *g*\* (·) represents the adjoint of *g*(·). Similarly, 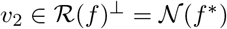

#### Remark 2.

Since the optimal solution of (4) is *u*_1_ = *g*(*υ*_1_), *g*(*υ*_2_) = 0 holds, i.e., 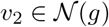 and thus *g* = *f* *. This also satisfies *f* (*u*_2_) = 0 since 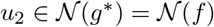.

#### Theorem 1

For a linear crosswise mapping case, the optimal crosswise mappings (*f, g*) for (4) are the adjoint to each other, i.e., *f* = *g*\* and *g* = *f*\*.

*Proof:* By Proposition 1 and Remark 1 and 2.

## III. Extension to deep learning model

Motivated by the theoretical derivation and result from the linear mapping case, we extend this conceptual framework to deep learning practice by utilizing a cross-domain autoencoding architecture and a separate gate layer to identify domain-specific information in a self-supervised manner. We also consider a cycle-consistency concept as free supervisory signal for learning a crosswise mapping (*f,g*) across two domains.

### A. XAE Learning Architecture

To address cross-domain translation in a deep learning setting, we consider crossed autoencoder (XAE) architecture as a pair of domain-specific autoencoder networks composed of encoders Φ_A_ and Φ_B_ and decoders Ψ_*A*_ and Ψ_*B*_ that independently learn latent representations of data from independent domains *A* and *B* in a joint latent space *z* as shown in Fig. 3. The self-supervised autoencoder system learns to minimize domain-specific autoencoder reconstruction loss while concurrently learning to minimize crossdomain cycle consistency loss following domain translation through the shared latent representation and performs crossdomain translation by swapping the decoder networks during evaluation such that *f*_*A*→*B*_(*A_i_*) = Ψ_*B*_ (Φ_*A*_(*A_i_*)) and *f*_*B*→*A*_(*B_i_*) = Ψ_*A*_(Φ_*B*_ (*B_i_*)).

**Fig. 3.**
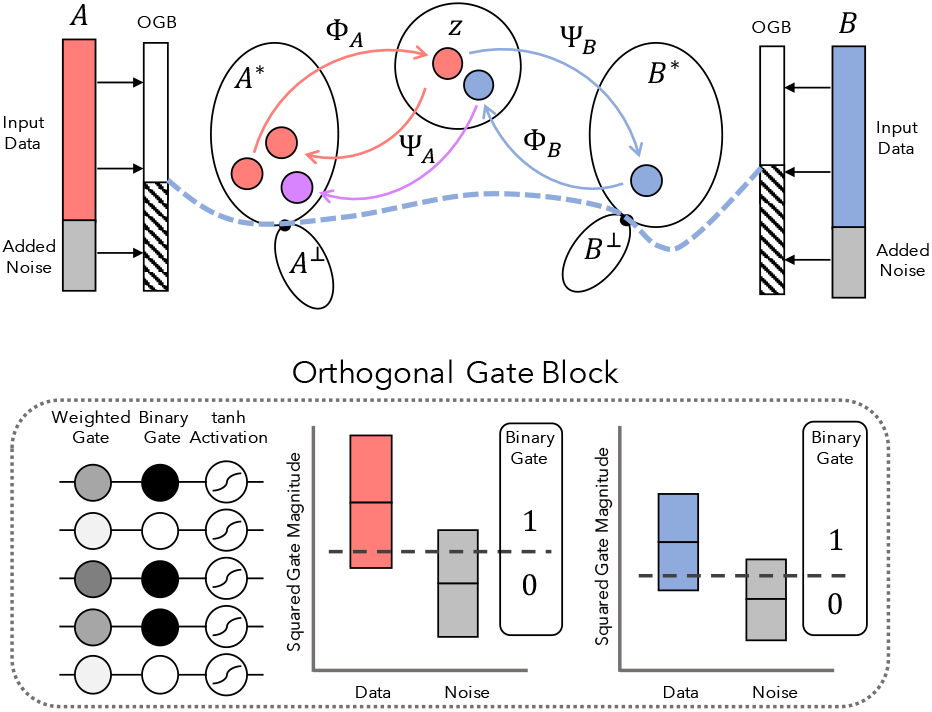
Conceptual illustration of architecture: we introduce an Orthogonal Gate Block to disentangle shared (*A*\*, *B*\*) and unshared (*A*^⊥^,*B*^⊥^) information in complementary domains during domain translation. Our approach seeks to minimize the cycle consistency loss between an input and its autoencoded reconstruction (red dot to red dot, in *A*, Eq. (5) and cycled reconstruction (red dot to purple dot in *A*, Eq. (6). Concordant latent representation is encouraged by minimizing cross-domain representations (red dot and blue dot in *z*, Eq. 7).

The learning objective for the XAE model includes terms to jointly optimize both domain-specific autoencoders as well as the bi-directional cycle consistent domain translation. Each domain-specific autoencoder learns to minimize a variational autoencoder objective which is composed of two terms in which the first term seeks to minimize the error between an input and its reconstruction while the second term seeks to minimize the Kullback-Leibler (KL) divergence between the distribution of learned latent features and an expected normal prior, such that

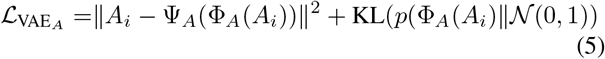

Further, the model learns to minimize domain cycle consistency by minimizing the reconstruction error of an input sample and its cycled reconstruction such that

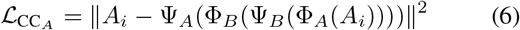

Lastly, because a single data point represented by either encoder should be represented by approximately the same latent representations, we introduce a novel mutual encoding divergence loss term to penalize mutual representational error such that

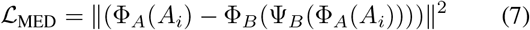

Lambda coefficients are chosen to approximate equally weighted loss term contributions, such that the full XAE learning objective is given by

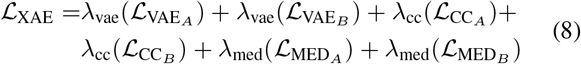

In this example the naive model assumes that the full information content provided in a given sample should be employed to generate cross-domain predictions, however in cases where this is not expected, we desire a mechanism to separate domain-specific from domain-shared information. Models are optimized with the Adam optimizer [9] with base learning rate of 1e–3 for 10 epochs.

### B. Orthogonal Gate Block

To consider autoencoder type mappings for *ϕ* and *ψ*, it is not straightforward to deal with dynamic changes of architectures for domain-specific and sharable content. To disentangle domain-specific information from information shared between domains, mutually informative data features are estimated through the introduction of a novel Orthogonal Gate Block (OGB). As shown in Figure 3, the OGB is composed of a single regularized element-wise multiplication layer 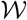, a non-trainable yet programmable binary elementwise multiplication layer 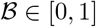, and a hyperbolic tangent activation function. The OGB appends noise sampled from the input domain’s respective data space to the input tensor and performs element-wise multiplication operations on an input data point *X* such that the layer’s function is defined as 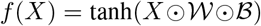 in which 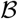 enforces orthogonality between separable and non-separable content.

Models are initiated with random weights, though after initial epochs of training, the magnitudes of the learned weights are used discriminate between meaningful data features and noise to enable cross-domain translation given an information bottleneck in *z*. First, features are randomly selected from the relevant data spaces and appended to the input data tensor. Then, after a fixed number of training epochs, the OGB evaluates the distribution of the gate weights associated with noise features and sets a threshold cutoff, which we set experimentally to be the 50% quantile of the noise-associated weight distribution, though this value is dependent on the question under inquery. Lastly, the OGB explicitly sets its binary layer to remove either data or noise features whose learned gate weight magnitude is lower than the established cutoff and in so doing enforces orthogonality onto the feature set that explicitly passes through relevant features while filtering out features that are learned to be lowly weighted in performing the cross-domain translation task.

#### Lemma 4

Magnitude-based Weight Pruning satisfies an orthonormal transformation condition.

*Proof :* (by definition) Magnitude-based weight pruning function result in sparse connection, a fully connected layer whose parameters’ values are either zero or 1 and thus, it only allows bijective connection. Then we can consider it as *W* where each row of *W*, *w_i_* has only one non-zero component and 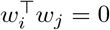 where *i* ≠ *j*.

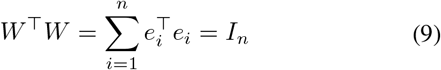

where *e_i_* represents the standard basis or unit vector of *n*-dim space.

## IV. Results and Discussion

To evaluate the efficacy of our proposed approach, we consider a simple dataset with perfectly overlapping mutual information which is modeled using the MNIST dataset of hand-written digits in 28×28 pixel image (2D) form, and a vectorized (1D) 784×1 pixel form. We perform a series of experiments by adding increasing volumes of noise to the original 784 pixel one-dimensional input tensor. We train several XAE models for 100 epochs on the two unpaired data sets described above. Figure 4.A illustrates the magnitude of learned weights in the orthogonal gate block at the end of training for five experiments each run with a different proportion of noise added to the data (0, 20%, 40%, 60%, and 80%). In all cases, we observe attenuation of noise features by their learned respective weights which may indicate that the OGB is learning to limit the importance of these features to the domain translation task. The squared magnitude of the learned weights serve as a reasonable discriminatory variable, as shown by the receiver operating characteristic curves in Figure 4.B, which we do not expect to perfectly discriminate between data and noise because several features of the input data such as the extreme perimeter of the MNIST images, are not expected to be important for domain translation.

**Fig. 4.**
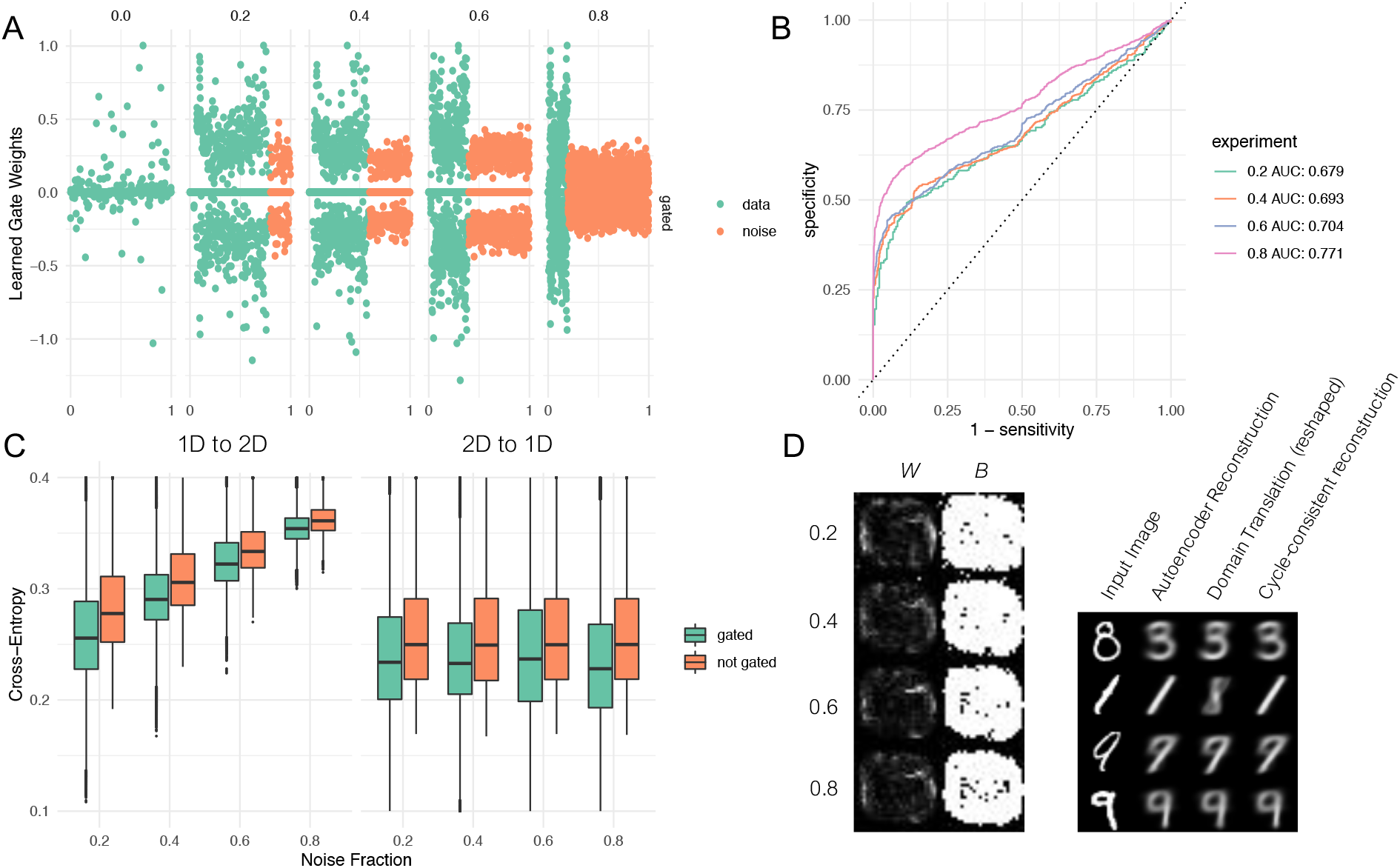
Evaluation of the XAE learning system. (A) learned gate weights of actual data (green) and injected noise (orange) along the long axis of vectorized images illustrates a tendency for the learning system to lowly weight features associated with noise. (B) ROC curves of data/noise separability strickly by learned gate weight. (C) Cross-entropy between an input in one domain and its prediction in the other illustrates an advance of utilizing the OGB in this particular domain translation task. (D) Illustration of learned gate weights and the learned corresponding binary layer illustrates that the OGB learned to identify features associated with the center of the images while ignoring the perimeter of the MNIST data set.

Because mutual information between the two domains is perfectly known, we fairly compare the domain translation prediction with knowable ground truth. Figure 4.C compares the cross-entropy between the predicted translations and their corresponding ground truth with and without the addition of the gate layer across experiments with varying noise burdens. Although slight, we observe a consistent improvement in the cross-entropy between prediction and ground truth with the addition of the OGB, suggesting that the layer’s removal of less-informative features facilitates better cross-domain prediction. In this example, the added noise was applied only to the 1D input data to illustrate a domain-specific affect of additive noise. Figure 4.D illustrate the resulting weights and binary layer of the OGB after training, which clearly illustrate that in general the OGB appears to automatically identify the center of the image as being relevant to the domain translation task. The accompanying figure illustrates four examples of a single input imaging and its various representations through the learning framework.

For visualization, we consider the original MNIST digit at the far left, followed by its direct autoencoded reconstruction, which is analogous to a typical variational autoencoder with characteristic softening of high-frequency detail. The third column illustrates the domain-translated sample, generated as a 1D vector but rearranged here to illustrate a comparable 2D representation, while the fourth column illustrates a cycle-consistent reconstruction of the input data point following domain translation. Note that these images are created by reconstructing a 2D image from the vectorized 1D OGB layer.

This work may be applicable to many computational tasks for which paired measurements from two complementary domains are absence, and in situations where measurements from either domain are expected to contain both domain-specific and sharable information content between the two. As deep learning methods continue to facilitate improved methods for domain translation, we anticipate a growing need for quantifying and identifying which features are mutually informative between complementary data domains. The approach presented herein is presently applied to a familiar benchmark dataset, but additional work is necessary to establish the generalizability of the principles set forth in this work.

With expected applications in the biomedical domain, we anticipate the need for synthetic biological data that accurately models unpaired domain-specific measurements with which to evaluate our approach. Future studies will incorporate additional synthetic data to further evaluate the capacity for this class of learning system to identify domain-specific features from complementary profiling technologies.

## V. Conclusion

In this work, we consider the domain translation problem in the absence of paired measurements. We motivate the need for identifying sharable and non-sharable information between complementary data domains when considering cross-domain translation. We introduce a neural network layer designed to separate domain-specific information when learning optimal domain translation models by appending noisy data features and utilizing the magnitude of feature-specific weighting to remove noise and non-informative features. On a benchmark dataset, we illustrate that the novel orthogonal gate block incorporated into a neural network learning architecture appears able to distinguish between sharable and non-sharable information in a purely data-driven manner.

## Acknowledgement

This work was supported in part by the National Cancer Institute (U54CA209988, U2CCA233280). YHC acknowledges the OHSU Center for Spatial Systems Biomedicine, Brenden Colson Center for Pancreatic Care and Biomedical Innovation Program Award from the Oregon Clinical & Translational Research Institute.

